# Modular, inducible, and titratable expression systems for *Escherichia coli* and *Acinetobacter baumannii*

**DOI:** 10.1101/2024.05.28.596346

**Authors:** Emily E. Bacon, Jennifer S. Tran, Nischala Nadig, Jason M. Peters

**Affiliations:** Pharmaceutical Sciences Division, School of Pharmacy, University of Wisconsin-Madison, Madison, WI 53705; Microbiology Doctoral Training Program, University of Wisconsin-Madison, Madison, WI 53706; Great Lakes Bioenergy Research Center, University of Wisconsin-Madison, Madison, WI 53726; Department of Bacteriology, University of Wisconsin-Madison, Madison, WI 53706; Department of Medical Microbiology and Immunology, University of Wisconsin-Madison, Madison, WI 53706; Center for Genomic Science Innovation, University of Wisconsin-Madison, Madison, WI 53706

**Keywords:** synthetic biology, gene expression, cloning, shuttle vector, Tn7 vector

## Abstract

Gene expression systems that transcend species barriers are needed for cross-species analysis of gene function. In particular, expression systems that can be utilized in both model and pathogenic bacteria underpin comparative functional approaches that inform conserved and variable features of bacterial physiology. Here, we develop replicative and integrative vectors alongside a novel, IPTG-inducible promoter that can be used in the model bacterium *Escherichia coli* K-12 as well as strains of the antibiotic-resistant pathogen, *Acinetobacter baumannii*. We generate modular vectors that transfer by conjugation at high efficiency and either replicate or integrate into the genome, depending on design. Embedded in these vectors, we also developed a synthetic, IPTG-inducible promoter, P_*abstBR*_, that induces to a high level, but is less leaky than the commonly used *trc* promoter. We show that P_*abstBR*_ is titratable at both the population and single cell level, regardless of species, highlighting the utility of our expression systems for cross-species functional studies. Finally, as a proof of principle, we use our integrating vector to develop a reporter for the *E. coli* envelope stress σ factor, RpoE, and deploy the reporter in *E. coli* and *A. baumannii*, finding that *A. baumannii* does not recognize RpoE-dependent promoters unless RpoE is heterologously expressed. We envision that these vector and promoter tools will be valuable for the community of researchers that study fundamental biology of *E. coli* and *A. baumannii*.

**Importance:** *Acinetobacter baumannii* is a multidrug-resistant, hospital-acquired pathogen with the ability to cause severe infections. Understanding the unique biology of this non-model bacterium may lead to the discovery of new weaknesses that can be targeted to treat antibiotic-resistant infections. Here, we provide expression tools that can be used to study gene function in *A. baumannii*, including in drug-resistant clinical isolates. These tools are also compatible with the model bacterium, *Escherichia coli*, enabling cross-species comparisons of gene function. We anticipate that the use of these tools by the scientific community will accelerate our understanding of *Acinetobacter* biology.

## Introduction

Historically, research in bacterial genetics focused on specific model organisms, such as *Escherichia coli* K-12, due to a lack of techniques, tools, reagents, genome sequences, and general knowledge of non-model bacteria (1, 2). As a result, much of our current understanding about the basic physiology of Gram-negative bacteria comes from *E. coli* (3, 4). Although most core cellular processes are likely conserved, gene function and regulation can vary subtly or even dramatically across species boundaries (4, 5). Such deviation is obvious in pathogens such as *Acinetobacter baumannii*, which has adopted many traits that are distinct from *E. coli* K-12—most notably extreme antibiotic resistance (6-8). With advances in DNA sequencing and synthesis as well as tools that democratize genetic analysis across species (e.g., CRISPR approaches (9)), there now exists an enormous opportunity to shrink the knowledge and technique gaps between model bacteria and clinically relevant pathogens. One simple approach to bridge the gap would be to develop systems capable of assessing gene function in both model and pathogenic bacteria, such that the function of any gene could be readily compared in different strain or species backgrounds.

Here, we focus on genetic tools that function in the antibiotic-resistant pathogen, *A. baumannii. A. baumannii* is considered an “urgent threat” by the Centers for Disease Control and Prevention due to its ability to resist nearly all available antibiotic treatments (10). Although some promising new anti-*Acinetobacter* compounds have recently been discovered (11, 12), more work is needed in this area as *Acinetobacter* is adept at acquiring and developing new resistance mechanisms (13-15). *A. baumannii* is poorly studied compared to *E. coli* K-12 and even other Gram-negative pathogens such as *Pseudomonas aeruginosa*; however, understanding the distinct physiology of *A. baumannii* is critical to developing new treatments (16, 17). For instance, lipid A, an essential component of the outer membrane in most Gram-negatives and a binding site for the antibiotic colistin (18), is not essential for viability in many *A. baumannii* strains including clinical isolates (19). Further, regulation of stress pathways that could play roles in antibiotic resistance, tolerance, or persistence is distinct in *A. baumannii* compared to other γ-proteobacteria, as *A. baumannii* lacks conserved transcription factors such as the stationary phase sigma (σ) factor, RpoS (20, 21).

Vectors that are capable of replicating in or integrating into *E. coli* and *A. baumannii* have been previously described, but also share important limitations. Replicative shuttle vectors typically use a high-copy, ColE1 origin of replication for *E. coli* and either the pWH1266 (22) or pRSF1010 (23) origin for *A. baumannii*. The pWH1266 and pRSF1010 origins are compatible in *A. baumannii*, enabling expression from two replicative vectors in the same cell (23). Integrative vectors based on the site-specific transposon Tn*7* insert DNA cargo into the genome downstream of the *glmS* gene and have been used extensively in *E. coli* (24), *A. baumannii* (25-27), and many other species (28, 29). However, many of these vectors were not designed to contain easily swappable modules (e.g., different antibiotic markers) outside of standard multiple cloning sites (MCS). Existing vectors typically employ inducible promoters that are either native to or designed for use in *E. coli* (30, 31). These include *E. coli* native promoters such as P_*l*_ac and P_*araBAD*_ that can be induced with IPTG or arabinose, respectively (22, 32), or semi-synthetic promoters such as P_*tac*_ and P_*trc*_ which are IPTG-inducible (23). Unfortunately, characteristics of these promoters pose challenges for precise control of expression. For instance, P_*araBAD*_ expression cannot be titrated with sub-saturating concentrations of its inducer, arabinose, due to “all or nothing” effects that result in a fraction of cells inducing at high level while others show minimal activity (33-35). P_*tac*_ and P_*trc*_ are sufficiently leaky that genes placed under their control often complement deletion phenotypes in the absence of inducer (30, 36, 37), and full induction often results in overexpression toxicity (38). A titratable promoter with less leakiness and a lower maximal level of expression would be ideal for physiological expression and gene function studies in *A. baumannii*.

In this work, we generate useful reagents for gene function studies in *A. baumannii* and *E. coli*. We create modular vectors that replicate or integrate in both species, and carry the novel promoter P_*abstBR*_, which can be induced and titrated with IPTG. In a proof of principle experiment, we combine all three reagents to probe the activity of the *E. coli* envelope stress σ factor, RpoE, in both species.

## Results and Discussion

### Modular replicative and integrating vectors for *E. coli* and *A. baumannii*

We sought to construct a modular set of replicative and integrative vectors that could be used to examine gene function in *A. baumannii* and *E. coli*. Our shuttle vector (Fig. 1a) replicates in *E. coli* using the medium copy origin, p15A (20-30 copies per cell (39)), and in *A. baumannii* using the low copy origin pWH1266 (∼9 copies per cell (40)). Our integrating vector (Fig. 1a) inserts into the genomes of *E. coli* and *A. baumannii* downstream of *glmS* using the Tn*7* transposase (provided on a separate plasmid (9, 28)). Both vectors have an antibiotic module flanked by XhoI sites for easily swapping resistance markers using Gibson assembly (41). Here, we have provided hygromycin, apramycin, and kanamycin versions of both replicative and integrative vectors. We note that hygromycin and apramycin are attractive resistance markers for studying multidrug-resistant pathogens given that neither antibiotic is used against *A. baumannii* clinically (25, 42). FRT sites in the integrative vector allow for optional FLP recombinase-mediated excision of the antibiotic marker (43, 44). The cloning module, or multiple cloning site (MCS), has several restriction sites for cloning genes of interest (Fig. 1b and 1c). Although other sites can be used, we recommend cloning into NcoI because it contains a translation start codon (ATG) in alignment with a strong upstream ribosome binding site (RBS) taken from the classic expression vector pTrc99a (45). The promoter module exists between AatII and NcoI sites for the replicating vector and SpeI and NcoI sites for the integrating vector. We provide these vectors with a novel, IPTG-inducible promoter (P_*abstBR*_, described below), but other promoters and RBSs of interest can be readily swapped into the module. Additionally, both the replicative and integrative vectors can be used in the same strain as multiple markers are available and only one vector replicates, ruling out compatibility issues.

**Figure 1.**
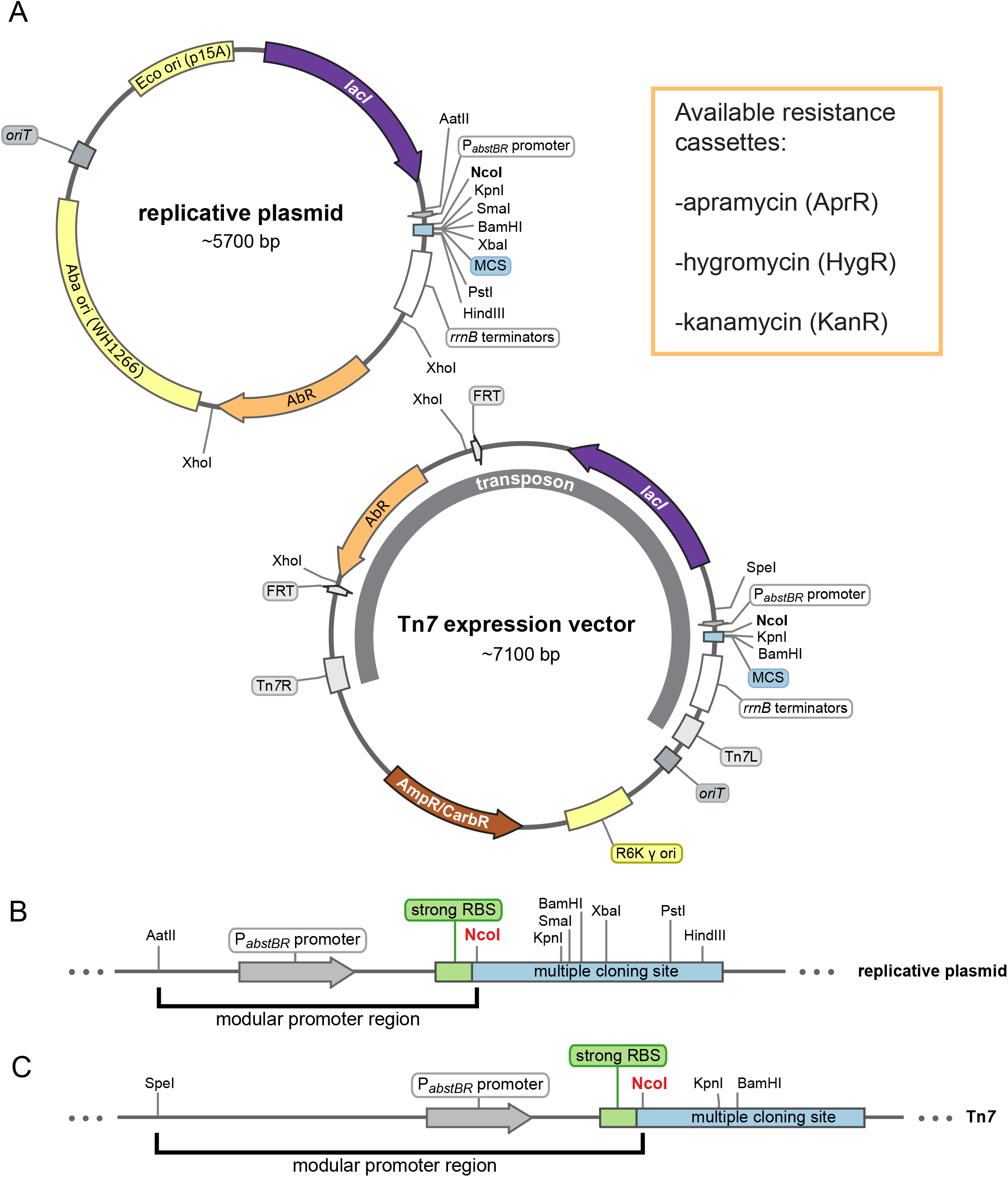
Modular replicative and integrative expression vectors. **(A)** Circular plasmid map and features of the replicative shuttle vector containing both *E. coli* and *A. baumannii* origins of replication (top) and the Tn*7* expression vector containing a transposon that will integrate into the chromosomal *att*_Tn*7*_ site (bottom). Available antibiotic resistance cassettes (AbR) are listed. Maps are adapted from SnapGene (GSL Biotech). **(B and C)** Linear maps showing the modular promoter region and multiple cloning sites (MCS) for the replicative plasmid and Tn*7* vector. NcoI site provides an ATG start codon optimally proximal to a strong ribosome binding site (RBS).

We next determined the efficiency of transfer for both vectors into *E. coli* and *A. baumannii*. Both vectors contain *oriT* sites, enabling transfer by conjugation from *E. coli* cells that are auxotrophic for diaminopimelic acid (DAP^-^) to DAP^+^recipient bacteria followed by antibiotic selection to recover only vector-containing recipients. Additionally, both vectors can be transferred by electroporation into competent recipient cells, if desired. To quantify efficiency of transfer by conjugation, we mated DAP^-^*E. coli* donor cells (*E. coli* K-12 WM6026) with model strains of *E. coli* K-12 (BW25113) and *A. baumannii* (ATCC 17978). We found that both vectors were transferred at efficiencies consistent with use in downstream experiments ranging in scale from individual genes to large libraries (Fig. S1a and S1b). Transfers of both the replicative and integrative vectors were highly efficient in *E. coli* (>10^-1^efficiencies for both vectors) and *A. baumannii* (>10^-2^ and 10^-4^efficiencies for replicative and integrative vectors, respectively). Importantly, our observed transfer efficiencies were on par with those needed for library construction for genome-scale experiments (9). We note that we observed instances of unintended integration of the Tn*7* vector backbone in both *E. coli* and *A. baumannii* (i.e., co-integrates (46)). The presence of such co-integrates in recipient colonies can be tested by screening for the *ampR*/*bla* gene (which confers carbenicillin resistance) present in the vector backbone. We patched 40 transconjugants for each organism, and while the frequency of integration with the vector backbone was relatively low (≤3/40 for each), we recommend testing transconjugants to verify insertion accuracy (Fig. S1c). Taken together, we have created modular replicative and integrative vectors for *E. coli* and *A. baumannii* that can be transferred at efficiencies that are useful for a variety of applications.

### A tightly regulated, IPTG-inducible promoter for *E. coli* and *A. baumannii*

We sought to develop an IPTG-inducible promoter with low leakiness and high expression for *A. baumannii*. We previously found that a broadly utilized synthetic promoter in *E. coli*, P_*LlacO-1*_, was unstable when used to express a toxic protein in *A. baumannii* (dCas9) (27). When we selected for mutants with stable expression of dCas9, we found that *lacO* repeats in the promoter had collapsed, creating a new IPTG-regulated promoter (Fig. 2a, *Acinetobacter* Suppressor of Toxicity or P_*abst*_). We hypothesized this promoter was weaker due to its success at repressing toxicity. To measure promoter activity in *A. baumannii*, we cloned P_*abst*_ upstream of a gene encoding Superfolder Green Fluorescent Protein (*sfgfp*) in our replicative vector (Fig. 2b). Our measurements confirmed that P_*abst*_ expression was very weak, with less than 2-fold increase in expression at saturating levels of inducer. This weak activity is likely due to divergence between the P_*abst*_ -35 element (TTATAA) and the consensus σ^70^-35 (TTGACA), especially at the -33 position (A versus G, respectively).

**Figure 2.**
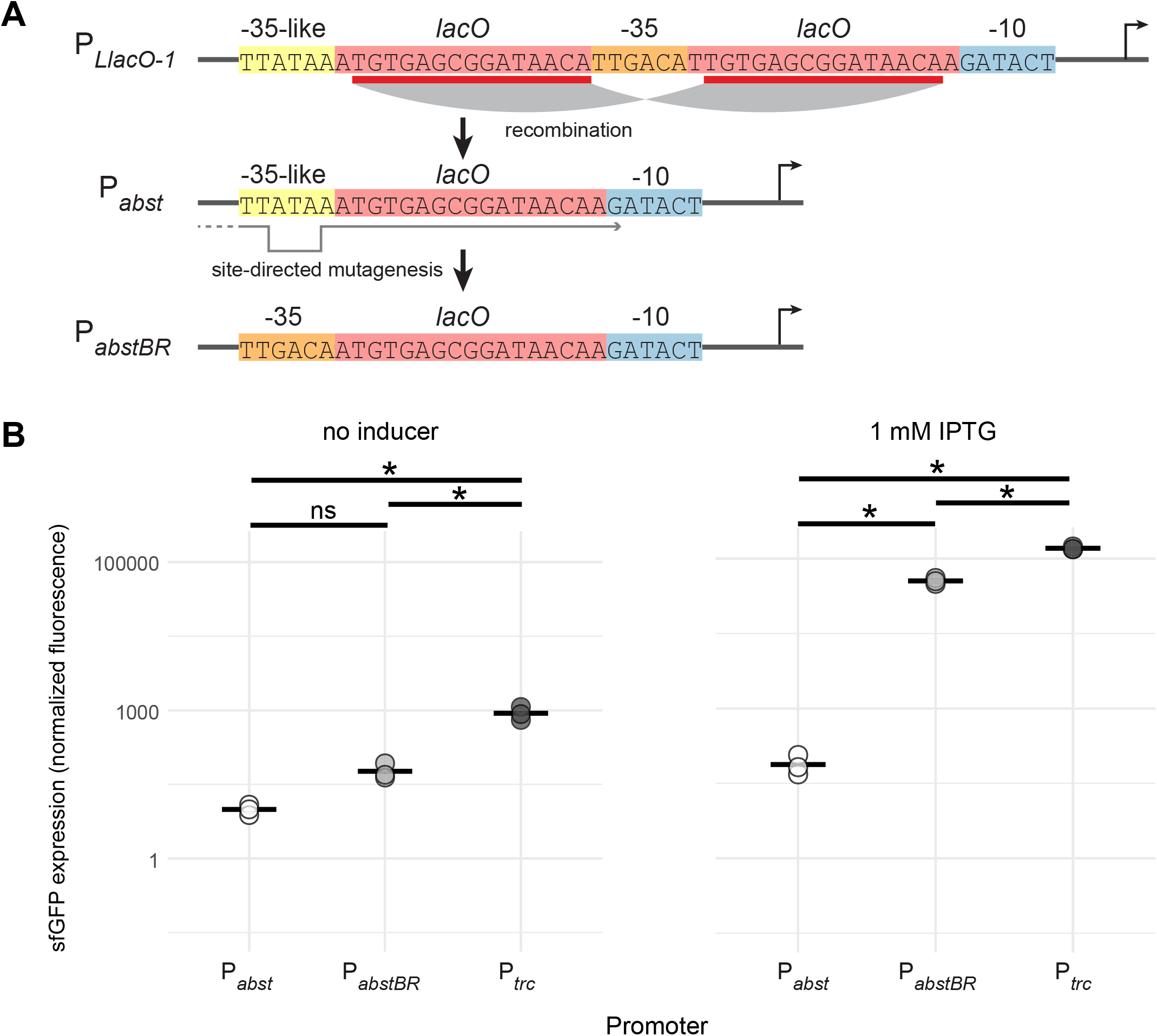
P_*abstBR*_ promoter construction and expression. **(A)** Promoter sequences showing the homologous recombination event in *lacO* repeat regions (red) of the P_*LlacO-1*_ sequence that produces P_*abst*_, which contains a -35-like region (yellow). Site-directed mutagenesis reverts the -35 region back to consensus (orange) to create P_*abstBR*_. **(B)** Dot plots showing sfGFP fluorescence from replicative vectors containing *sfgfp* under P_*abst*_, P_*abstBR*_, or P_*trc*_ promoters in *A. baumannii* ATCC 17978 with no IPTG (left) or 1 mM IPTG (right). Values were normalized to empty vector controls, and sample means are represented by a solid horizontal line (n=3). Asterisks and ns indicate significant and not significant sample differences, respectively (Welch’s *t-*tests; p-values < 0.05).

To generate a new promoter with higher activity but without repeating *lacO* elements, we used site-directed mutagenesis to replace the P_*abst*_ -35 sequence with a consensus -35 (Fig. 2a). We found that the new promoter, P_*abstBR*_ (*Acinetobacter* Suppressor of Toxicity with Better Regulation), showed significantly higher induction than P_*abst*_ (∼150-fold; Welch’s *t*-test, p=0.003) in *A. baumannii* (Fig. 2b). P_*abstBR*_ also showed ∼3-fold reduced leakiness compared to P_*trc*_, a popular IPTG-inducible promoter used in both *E. coli* (30) and *A. baumannii* (23); although induction at saturating levels of IPTG was somewhat lower (∼3-fold) than P_*trc*_. With reduced leakiness and a more physiologically appropriate expression range, P_*abstBR*_ has advantages for complementation and expression with reduced toxicity (36, 47).

### P_*abstBR*_ expression is titratable at the population and single cell level

Investigators frequently titrate promoter activity to determine expression-phenotype relationships and avoid toxic overexpression. To determine if P_*abstBR*_ expression is titratable at the population level, we induced expression of P_*abstBR*_-*sfgfp* at varying concentrations of IPTG from both our replicative and integrative vectors in *E. coli* K-12 BW25113 and *A. baumannii* ATCC 17978 (Fig. 3a and 3b). We found that P_*abstBR*_ was titratable in all tested contexts. Plasmid-borne P_*abstBR*_ showed similar patterns of IPTG induction in both *E. coli* and *A. baumannii* and had ∼10-fold higher level of maximal expression compared to an integrated copy. Unexpectedly, Tn*7* integrated P_*abstBR*_ showed a higher apparent level of expression in *A. baumannii* compared to *E. coli* at nearly every concentration of IPTG, including saturating concentrations (Fig. 3b). In addition to 17978, the *A. baumannii* field uses strains ATCC 19606 and AB5075 as antibiotic susceptible and resistant models, respectively. To test P_*abstBR*_ titratability in those strain backgrounds, we again expressed P_*abstBR*_-*sfgfp* at varying IPTG concentrations (Fig. S3). As expected, we found that P_*abstBR*_ was titratable at the population level.

**Figure 3.**
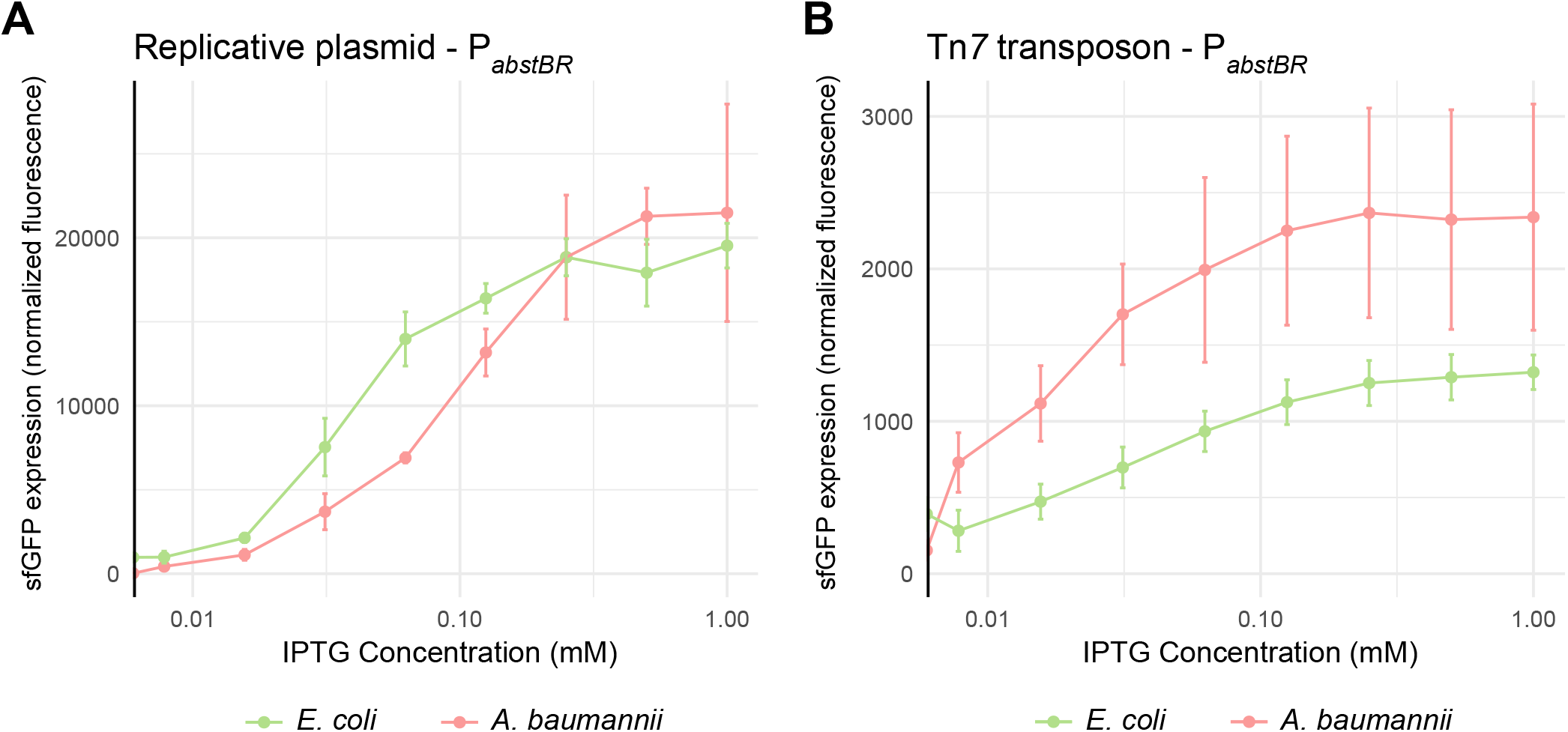
Titration of P_*abstBR*_ expression at the population level. Titration of expression from **(A)** the replicative plasmid or **(B)** the Tn*7* transposon. Plots shown are normalized sfGFP levels expressed from P_*abstBR*_ across IPTG concentrations for *E. coli* BW25113 and *A. baumannii* ATCC 17978. Error bars represent standard deviation (n=3 for replicative vector, n=6 for Tn*7* transposon).

Inducible promoters can erroneously appear to be titratable at the population level due to varying subpopulations of fully induced cells, as is seen in systems with active transport and feedback of inducer molecules (e.g., arabinose and P_*araBAD*_ (33)). To rule out this possibility, we measured induction of P_*abstBR*_-*sfgfp* at varying concentrations of IPTG in single cells using flow cytometry (Fig. 4a and 4b). We measured P_*abstBR*_ expression from replicative vectors as we reasoned that variations in plasmid copy number would be more likely to have a subpopulation effect. We found that P_*abstBR*_ was fully titratable at the single cell level in *E. coli* K-12 BW25113 and *A. baumannii* ATCC 17978. Distributions of sfGFP fluorescence were unimodal at all IPTG concentrations in both species, consistent with relatively uniform induction of P_*abstBR*_ at the single cell level. Although increasing concentrations of IPTG fully shifted the sfGFP distributions in *A. baumannii*, the distributions were wider than those seen in *E. coli* for unknown reasons (Fig. 4b). One possibility to explain increased expression variation in *A. baumannii* is simply that the pWH1266 origin has intrinsically greater plasmid copy number variation than p15A, although testing plasmid copy number at the single cell level is fraught with challenges (48). We conclude that P_*abstBR*_ is titratable at the single cell level, enabling gene function studies with precise levels of expression.

**Figure 4.**
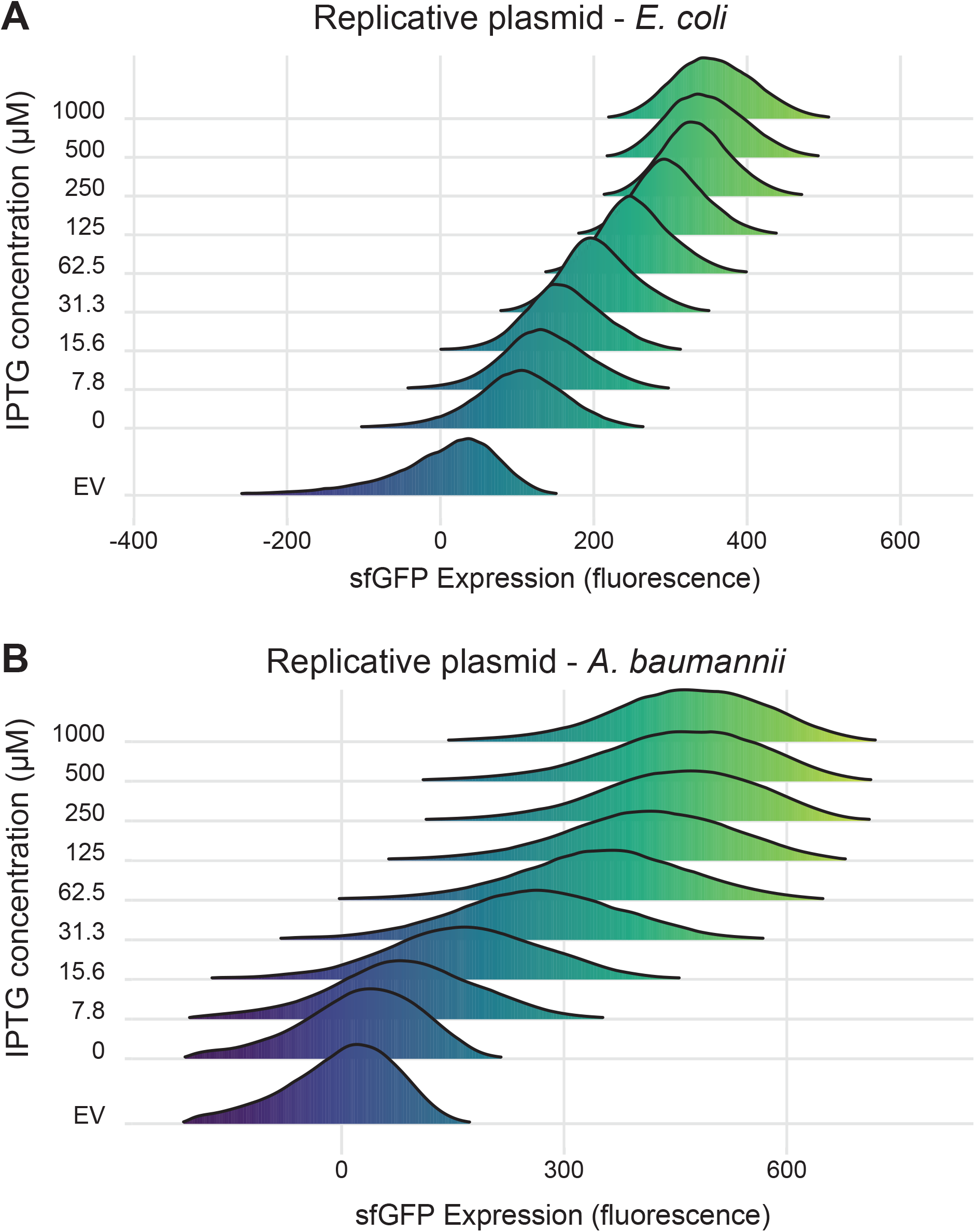
Titration of P_*abstBR*_ expression at the single-cell level. Titration of expression in **(A)** *E. coli* BW25113 or **(B)** *A. baumannii* ATCC 17978. Ridgeline plots depict overlapping density plots of sfGFP fluorescence for cells induced at different IPTG concentrations, measured by flow cytometry and expressed from the replicative expression vector under control of P_*abstBR*_. EV are empty vector (no GFP) control samples in 1 mM IPTG.

### Modular vectors and P_*abstBR*_ enable gene regulation studies in *E. coli* and *A. baumannii*

As a proof of principle to demonstrate the utility of our P_*abstBR*_ vector set in studying gene function, we investigated RpoE promoter activity in *E. coli* and *A. baumannii*. RpoE, also known as σ^E^, is an extracytoplasmic function (ECF) σ factor that regulates the envelope stress response in *E. coli* and related γ-proteobacteria (49-52). Species as distant from *E. coli* as *Pseudomonas aeruginosa* have a functional ortholog (AlgU, 66% identity) that recognizes the same DNA sequence as RpoE (53); however, a BLAST search of the *A. baumannii* genome recovered no hits for RpoE. To determine if *A. baumannii* recognizes RpoE-dependent promoters, we cloned the autoregulated *rpoE* promoter (P_*rpoE*_) from *E. coli* into our integration vector upstream of a gene encoding monomeric Red Fluorescent Protein (*mrfp*) as a reporter. We integrated this construct into both *E. coli* and *A. baumannii* and found P_*rpoE*_ was only active in *E. coli* (Fig 5a and 5b). To determine if the promoter could be recognized in *A. baumannii* in the presence of RpoE, we cloned the *rpoE* gene into our replicating vector under the control of P_*abstBR*_. We found that expression of RpoE in *A. baumannii* was sufficient to drive expression from P_*rpoE*_ (Fig 5a). This suggests that *A. baumannii* has no RpoE activity and that no other factors in *A. baumannii* can recognize RpoE promoters. As expected, we also found that overexpression of RpoE in *E. coli* resulted in increased P_*rpoE*_ activity (Fig. 5b). Importantly, these results demonstrate the ability to utilize our integrative and replicative expression systems together, in the same strain, to better understand biology and gene function in both *E. coli* and *A. baumannii*.

**Figure 5.**
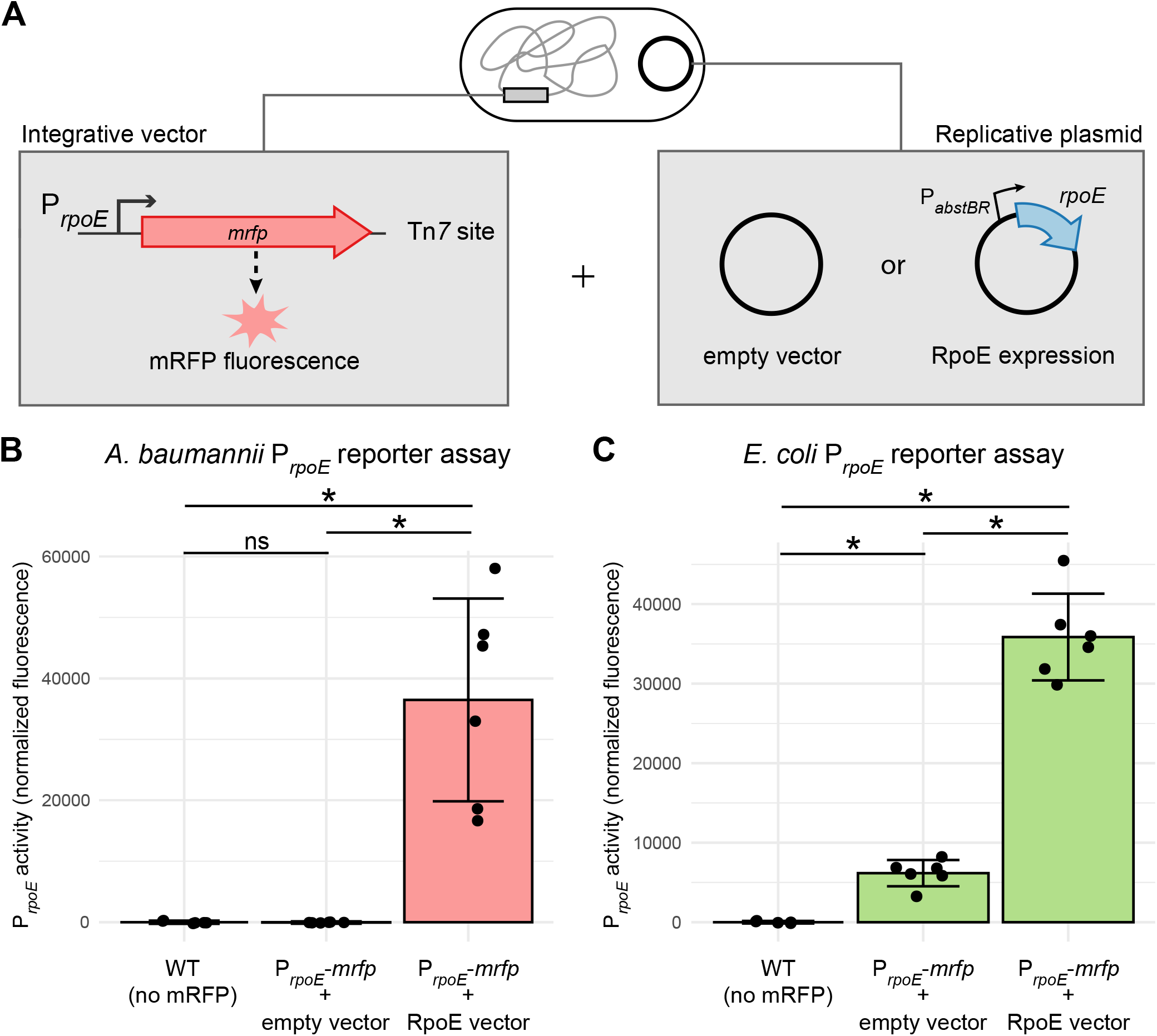
Modular integrative and replicative vectors facilitate a functional reporter assay. **(A)** Graphical depiction of reporter assay experiments. Strains contain an mRFP reporter under control of the *E. coli*-native *rpoE* promoter (P_*rpoE*_) in the *att*_Tn*7*_ site (constructed using the Tn*7* vector) and either a P_*abstBR*_-*rpoE* overexpression vector or empty vector control (replicative plasmid). **(B and C)** Bar graphs of mRFP fluorescence from P_*rpoE*_ with and without expression of RpoE *in trans* from the replicative plasmid in *E. coli* or *A. baumannii*. As RpoE is native to *E. coli*, the *E. coli* strains also carry a copy of the *rpoE* gene on the chromosome. Fluorescence is normalized to no mRFP controls, and individual data points and standard deviation are displayed (n=6). Asterisks and ns indicate significant and not significant sample differences, respectively (Welch’s *t*-tests; p-values < 0.05)

### Conclusion

Here, we have provided modular vectors that replicate and integrate into *E. coli* and *A. baumannii*, and a titratable, IPTG-inducible promoter, P_*abstBR*_. We envision that our vectors will be valuable for complementation studies, particularly for comparing the function of genes in *E. coli* to those found in *A. baumannii*. We predict that our tools will allow for precise tuning of gene expression to achieve physiological or somewhat higher levels of expression while avoiding toxicity from extreme high-level overexpression. As such, our vectors could also be used for expressing gene fusions with fluorescent proteins for localization studies. The high integration efficiencies make library scale experiments possible, as we have previously shown for Tn*7*-based CRISPRi work (9). Given the host ranges of our vector components, we expect our vectors to be broadly useful for gene function studies in *Acinetobacter* species not tested here, including multidrug-resistant isolates.

## Materials and Methods

### Strains and growth conditions

Strains are listed in Table S1. *Escherichia coli* and *Acinetobacter baumannii* were grown in Lennox lysogeny broth (LB) at 37°C shaking in a flask at 250 rpm, in a culture tube on a rollerdrum at max speed, in a 96-well plate shaking at 900 rpm, or in a plate reader shaking (Tecan Infinite Mplex or Tecan Sunrise). Culture medium was solidified with 1.5% agar for growth on plates. Antibiotics were added when necessary: 100 μg/mL ampicillin (amp), 30 μg/mL kanamycin (kan), 50 μg/mL apramycin (apr), and 150 μg/mL hygromycin (hyg) for *E. coli* and 150 μg/mL carbenicillin (carb), 60 μg/mL kanamycin (kan), 100 μg/mL apramycin (apr), 150 μg/mL hygromycin (hyg) for *A. baumannii*. Diaminopimelic acid (DAP) was added at 300 μM to support growth of *E. coli* dap^-^donor strains. IPTG (isopropyl b-D-1-thiogalactopyranoside) was added at varying concentrations from 0 to 1 mM as indicated in the figures or figure legends. Strains were preserved in 15% glycerol at -80°C. Plasmids were propagated in *E. coli* strain BW25141 *att*_Tn*7*_::acrIIA4 (sJMP3053) or in *E. coli* strain DH10B (sJMP1) for DNA extraction and analysis or in *E. coli* strain WM6026 *att*_Tn*7*_::acrIIA4 (sJMP3257) for conjugation.

### General molecular biology techniques

A complete list of plasmids and oligonucleotides are listed in Tables S2 and S3. Oligonucleotides were synthesized by Integrated DNA Technologies (Coralville, IA). Plasmid DNA was purified using GeneJet Plasmid Miniprep kit (Thermo) or the Purelink HiPure Plasmid Midiprep kit (Invitrogen K210005). PCR was performed according to manufacturer directions using Q5, OneTaq, or Phusion DNA Polymerases (NEB). DNA was digested with restriction enzymes from NEB. PCR products were purified with DNA Spin and Concentrate kit (Zymo Research) following manufacturer instructions or gel-purified from kit (Zymo Research). Plasmids were assembled using NEBuilder HiFi DNA assembly kit (NEB). DNA was quantified on a Nanodrop Lite or Qubit. Plasmids and recombinant strains were sequenced via Sanger sequencing by Functional Biosciences or Oxford Nanopore sequencing by Plasmidsaurus.

### Construction of replicative expression vectors

Details for construction of expression vectors are listed under “Construction/notes” for corresponding vectors (Table S2). Briefly, base replicative expression plasmid construction was performed using HiFi assembly with: (i) p15A origin of replication and *oriT* from pJMP3262, (ii) pWH1266 origin of replication from pJMP3347, (iii) pTrc99a plasmid base including *lacI* and MCS from pJMP3067, and (iv) *kanR* marker from pJMP3341 to create plasmid pJMP3649. To swap the promoters, pJMP3649 was cut with AatII and NcoI enzymes and HiFi assembled with gblocks containing the desired promoters, to create plasmids pJMP3651 (P_*abst*_, *kanR*) and pJMP3653 (P_*abstBR*_, *kanR*). To swap the resistance markers, pJMP3653 was cut with XhoI enzyme and HiFi assembled with gblocks containing the desired resistance markers, to create plasmids pJMP3664 (P_*abstBR*_, *aprR*) and pJMP3665 (P_*abstBR*_, *hygR*). To test expression of genes from these vectors, the *kanR* versions of the vectors with P_*trc*_, P_*abst*_, and P_*abstBR*_ (pJMP3649, pJMP3651, and pJMP3653, respectively) were cut with NcoI and BamHI enzymes and HiFi assembled with the *sfgfp* gene amplified from pJMP2748 to create plasmids pJMP3650, pJMP3652, and pJMP3654.

### Construction of P_*abstBR*_

Site-directed mutagenesis of the P_*abst*_ promoter was performed by single-primer high-fidelity Phusion PCR using pJMP3407 and oJMP2167. The PCR product was treated with DpnI, electroporated into sJMP3053, and selected on kan to make plasmid pJMP4481 containing the P_*abstBR*_ promoter. The mutation was verified by whole-plasmid sequencing with Plasmidsaurus.

### Conjugative-based transfer of expression vectors

*Replicative vector:* Donor Dap^-^*E. coli* mating strain containing desired replicative expression vector and recipient strain (*A. baumannii* or *E. coli*) were both scraped off an agar plate into LB at OD600 of ∼3. Strains were mixed at equal ratios, placed on a 0.45 μm filter on an LB plate, and incubated upright at 37°C for ∼3 hrs. Filters were vortexed in LB media to remove cells and plated onto LB plates supplemented with appropriate antibiotic.

*Tn7 integrating vector:* Conjugation was performed similarly to above, except with the addition of a donor Dap^-^*E. coli* strain carrying a Tn*7* transposase plasmid (tri-parental mating) for *E. coli, A. baumannii* ATCC 17978, and AB5075 strains. For *A. baumannii* ATCC 19606, quad-parental mating was performed, using an additional Dap^-^donor *E. coli* strain (sJMP4061) harboring a helper plasmid that contains extra mating machinery to improve efficiency. Tn*7* matings were performed for ∼4 hrs before plating on LB plates supplemented with appropriate antibiotic.

Ten-fold serial dilutions were spotted (10 μL) on LB and LB with antibiotic. Transfer efficiencies were calculated as transformants or transconjugants (colony forming units or CFUs on selective plates) divided by total cells (CFUs on LB only).

### Promoter activity assays

Promoter activities were assayed using the sfGFP expression vectors. Promoter-*sfgfp* or empty vector strains were grown to saturation in LB supplemented with appropriate antibiotic and IPTG inducer, washed several times with 1xPBS to remove all media, and GFP fluorescence and OD_600_ were measured in a Tecan Infinite Mplex plate reader. Values were normalized to OD_600_ readings and were background-subtracted using empty vector cells.

### Flow cytometry

Cells containing either a P_*abstBR*_-*sfgfp* vector or empty vector control were grown in LB supplemented with kan and varying concentrations of IPTG to saturation overnight in tubes. Cells were formaldehyde fixed, washed, and resuspended in 1xPBS. GFP fluorescence was measured by flow cytometry on a LSR Fortessa instrument (BD Biosciences) at 100,000 events/sample. Data were analyzed in FlowJo (FlowJo, LLC) using singlet gates and dead cell or debris exclusion gates, as previously described (54).

## Data availability

Plasmids and their sequences are available from Addgene under accession numbers xxxx-xxxx (note: accession #s pending). R code for data analysis and graphs can be found at https://github.com/jasonpeterslab/Aba-Eco-expression-systems-2024. Data available on request.

## Acknowledgements

We thank Colin Manoil for providing AB5075 WT strain and Quanjiang Ji for pSGAb-km (Addgene plasmid # 121999). We also thank the UWCCC Flow Cytometry lab for equipment access and assistance (NIH Special BD LSR Fortessa Project: 1S100OD018202-01). This work was supported by the National Institutes of Health under award numbers K22AI137122 and 1R35GM150487-01. J.S.T. was funded by an NSF GRFP and the SciMed Graduate Research Scholars program.

