## Supplemental Material for "Modular, inducible, and titratable expression systems for *Escherichia coli* and *Acinetobacter baumannii*"

**Figure S1 Conjugation and transposition efficiencies. (A)** Dot plot of conjugation efficiency of the replicative vector to the recipient strain (*A. baumannii* ATCC 17978 or *E. coli* BW25113) after ~3-hour incubations. Bars represent mean efficiencies (n=3). Dashed line is drawn at the limit of detection, defined by the maximum number of colony-forming units (CFUs) observed on non-selective plates in each experiment. **(B)** Dot plot showing transposition efficiency of the Tn*7* transposon into the $att_{\mathrm{Tn7}}$ site of *A. baumannii* ATCC 17978 or *E. coli* BW25113 after ~4-hour incubation with donor strains, one carrying the Tn*7* transposon vector and another carrying the Tn*7* transposase vector. Bars represent mean efficiencies (n=3); dashed line shows limit of detection. **(C)** Table showing instances of carbenicillin-resistant (CarbR) transconjugants of 40 isolates tested for Tn*7* co-integrates in *A. baumannii* and *E. coli*.

**Figure S2 Titration of $P_{abstBR}$ expression.** Expression from **(A)** the replicative plasmid vector or **(B)** the Tn*7* transposon. Plots shown are normalized sfGFP levels expressed from $P_{abstBR}$ across IPTG concentrations for *A. baumannii* ATCC 19606 and *A. baumannii* AB5075. Error bars represent standard deviation (n=3 for replicative vector, n=6 for Tn*7* transposon).

**S1**

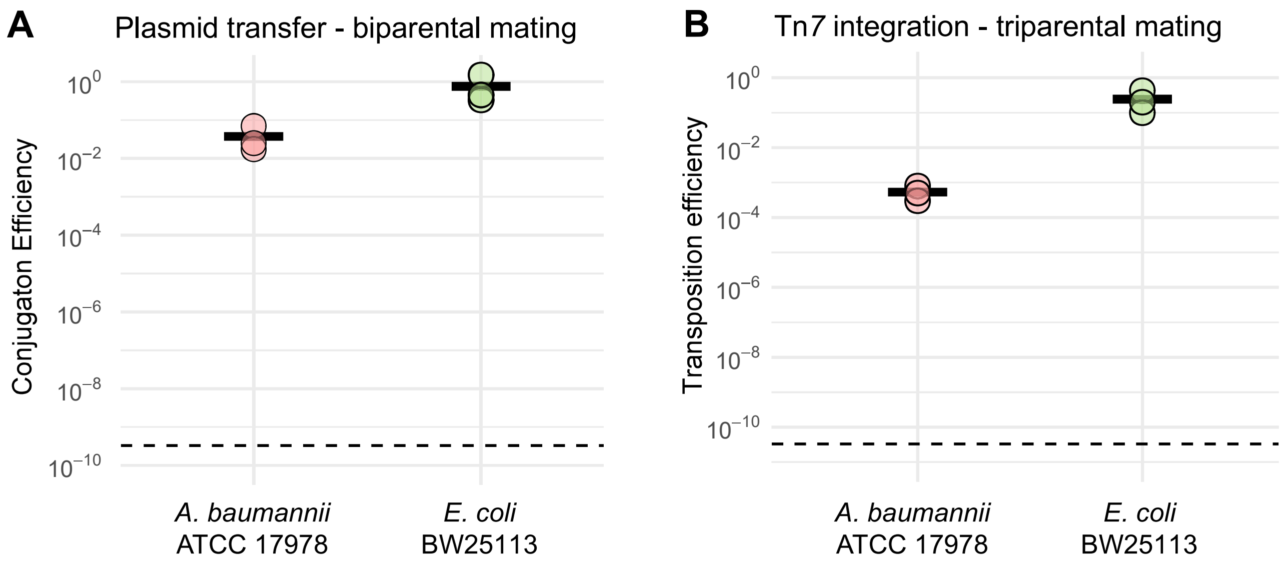


**C**

**Co-integrates: CarbR Tn*7* transconjugants**

| | Strain | |
|---|---|---|
| | *A. baumannii* ATCC 17978 | *E. coli* BW25113 |
| Tn*7* vector - empty | 2/40 | 0/40 |
| Tn*7* vector - sfGFP | 3/40 | 1/40 |

**S2**

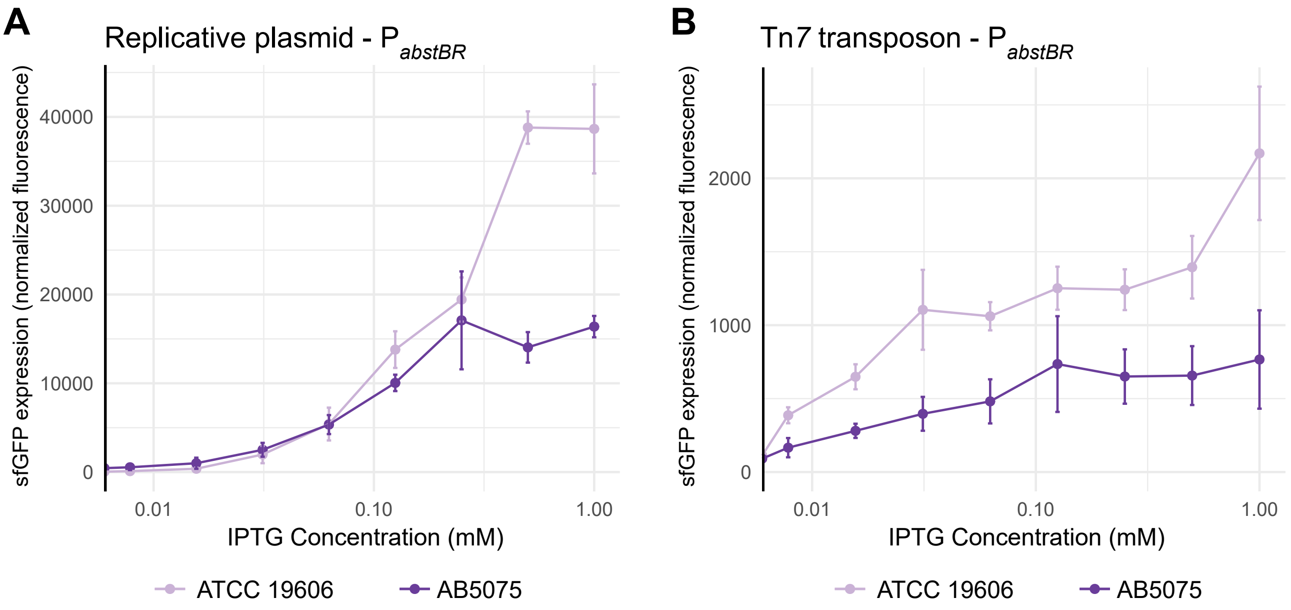
A
Replicative plasmid - P abstBR
sfGFP expression (normalized fluorescence)
40000
30000
20000
10000
0
0.01
0.10
1.00
IPTG Concentration (mM)
ATCC 19606
AB5075
B
Tn7 transposon - P abstBR
sfGFP expression (normalized fluorescence)
2000
1000
0
0.01
0.10
1.00
IPTG Concentration (mM)
ATCC 19606
AB5075

**Supplemental Table S1 Strains**

**Key**

| **Term** | **Description** |
|---|---|
| Strain # | JMP lab accession number |
| Description | Descriptive name, genotype, plasmids, construction details, phenotype, usage, etc |
| Source | Strains produced in this study unless otherwise noted |

| Strain # | Description[a] | Source |
|---|---|---|
| sJMP3053 | *Escherichia coli* (derived from BW25141) pir+, recAI cloning strain with anti-CRISPR (attTn7::acrIIA4); Δ*(araD-araB)*567, Δ*lacZ4787* (::*rrnB-3*), Δ(*phoB-phoR*)580, l-, *galU95*, Δ*uidA3*::pir+, recA1, endA9(del-ins)::FRT, *rph-1*, Δ(*rhaD-rhaB*)*568*, *hsdR514, attTn7::acrIIA4* | Ward, PMID: 38126769 |
| sJMP3075 | *Escherichia coli* MG1655 wild-type (sJMP163, CAG80011) | Carol Gross, UCSF |
| sJMP3076 | *Escherichia coli* BW25113 (sJMP6, CAG74538); F-, Δ(araD-araB)567, ΔlacZ4787(::rrnB-3), λ-, rph-1, Δ(rhaD-rhaB)568, hsdR514 | Carol Gross, UCSF |
| sJMP3257 | *Escherichia coli* (derived from WM6026) pir+, dap- mating strain with anti-CRISPR (attTn7::acrIIA4); lacIq, rrnB3, DElacZ4787, hsdR514, DE(araBAD)567, DE(rhaBAD)568, rph-1 att-lambda::pAE12-del (oriR6K/cat::frt5), del 4229(dapA)::frt(DAP-), del(endA)::frt, uidA(delMluI)::pir(wt), attHK::pJK1006::del1/2(del oriR6K-cat::frt5, del trfA::frt), attTn7::acrIIA4 | Banta, PMID: 38496613 |
| sJMP3261 | *Escherichia coli* sJMP3257 mating strain with plasmid pTn7C1 (pJMP1039) transposase expression; ampR, dap- | Banta, PMID: 38496613 |
| sJMP3329 | *Acinetobacter baumannii* ATCC 19606 WT | ATCC |
| sJMP3348 | *Acinetobacter baumannii* ATCC 17978 WT | ATCC |
| sJMP3518 | *Acinetobacter baumannii* AB5075 WT | Colin Manoil, UW |
| sJMP3683 | *E. coli* sJMP3257 mating strain with plasmid pJMP3649 (Ptrc EV); kanR, dap- | this study |
| sJMP3684 | *E. coli* sJMP3257 mating strain with plasmid pJMP3650 (Ptrc sfGFP); kanR, dap- | this study |
| sJMP3685 | *E. coli* sJMP3257 mating strain with plasmid pJMP3651 (Pabst EV); kanR, dap- | this study |
| sJMP3686 | *E. coli* sJMP3257 mating strain with plasmid pJMP3652 (Pabst sfGFP); kanR, dap- | this study |
| sJMP3687 | *E. coli* sJMP3257 mating strain with plasmid pJMP3653 (PabstBR EV kan); kanR, dap- | this study |
| sJMP3688 | *E. coli* sJMP3257 mating strain with plasmid pJMP3654 (PabstBR sfGFP kan); kanR, dap- | this study |
| sJMP3689 | *E. coli* sJMP3257 mating strain with plasmid pJMP3664 (PabstBR EV apr); aprR, dap- | this study |
| sJMP3690 | *E. coli* sJMP3257 mating strain with plasmid pJMP3666 (PabstBR sfGFP apr); aprR, dap- | this study |
| sJMP3691 | *E. coli* sJMP3257 mating strain with plasmid pJMP3665 (PabstBR EV hyg); hygR, dap- | this study |
| sJMP3692 | *E. coli* sJMP3257 mating strain with plasmid pJMP3667 (PabstBR sfGFP hyg); hygR, dap- | this study |
| sJMP3715 | *A. baumannii* 17978 (sJMP3348) with plasmid pJMP3649 (Ptrc EV); kanR | this study |
| sJMP3716 | *A. baumannii* 17978 (sJMP3348) with plasmid pJMP3650 (Ptrc sfGFP); kanR | this study |
| sJMP3717 | *A. baumannii* 17978 (sJMP3348) with plasmid pJMP3651 (Pabst EV); kanR | this study |
| sJMP3718 | *A. baumannii* 17978 (sJMP3348) with plasmid pJMP3652 (Pabst sfGFP); kanR | this study |
| sJMP3719 | *A. baumannii* 17978 (sJMP3348) with plasmid pJMP3653 (PabstBR EV); kanR | this study |
| sJMP3720 | *A. baumannii* 17978 (sJMP3348) with plasmid pJMP3654 (PabstBR sfGFP); kanR | this study |
| sJMP3721 | *A. baumannii* 19606 (sJMP3329) with plasmid pJMP3664 (PabstBR EV); aprR | this study |
| sJMP3722 | *A. baumannii* 19606 (sJMP3329) with plasmid pJMP3666 (PabstBR sfGFP); aprR | this study |
| sJMP3725 | *A. baumannii* AB5075 (sJMP3518) with plasmid pJMP3665 (PabstBR EV); hygR | this study |
| sJMP3726 | *A. baumannii* AB5075 (sJMP3518) with plasmid pJMP3667 (PabstBR sfGFP); hygR | this study |
| sJMP3733 | *E. coli* BW25113 (sJMP3076) with plasmid pJMP3653 (PabstBR EV); kanR | this study |
| sJMP3734 | *E. coli* BW25113 (sJMP3076) with plasmid pJMP3654 (PabstBR sfGFP); kanR | this study |
| sJMP3844 | *A. baumannii* 17978 (sJMP3348) with attTn7::PrpoE-mRFP reporter (Tn7 from pJMP3748; rpoE promoter sequence from E. coli), aprR | this study |
| sJMP3845 | *E. coli* (sJMP3075) with attTn7::PrpoE-mRFP reporter (Tn7 from pJMP3748; rpoE promoter sequence from E. coli), aprR | this study |
| sJMP4061 | *E. coli* mating strain with helper plasmid pEVS104 (sJMP2935; helper strain); kanR, dap- | Ward, PMID: 38126769 |
| sJMP12051 | *A. baumannii* 17978 (sJMP3348) with attTn7::PabstBR-sfGFP (Tn7 from pJMP12049); aprR | this study |
| sJMP12053 | *A. baumannii* 17978 (sJMP3348) with attTn7::PabstBR-empty (Tn7 from pJMP12042); aprR | this study |
| sJMP12055 | *A. baumannii* 19606 (sJMP3329) with attTn7::PabstBR-sfGFP (Tn7 from pJMP12049); aprR | this study |
| sJMP12057 | *A. baumannii* 19606 (sJMP3329) with attTn7::PabstBR-empty (Tn7 from pJMP12042); aprR | this study |
| sJMP12059 | *E. coli* BW25113 (sJMP3076) with attTn7::PabstBR-sfGFP (Tn7 from pJMP12049); aprR | this study |
| sJMP12061 | *E. coli* BW25113 (sJMP3076) with attTn7::PabstBR-empty (Tn7 from pJMP12042); aprR | this study |
| sJMP12063 | *E. coli* sJMP3257 mating strain with plasmid pJMP12042 (Tn7 PabstBR EV); ampR, aprR, dap- | this study |
| sJMP12065 | *E. coli* sJMP3257 mating strain with plasmid pJMP12049 (Tn7 PabstBR sfGFP); ampR, aprR, dap- | this study |
| sJMP12070 | *E. coli* mating strain with plasmid pJMP3853 (PabstBR RpoE OE vector); kanR, dap- | this study |
| sJMP12074 | (pJMP3853), aprR, kanR | this study |
| sJMP12076 | kanR | this study |
| sJMP12083 | *A. baumannii* AB5075 (sJMP3518) with attTn7::PabstBR-empty (Tn7 from pJMP12042); aprR | this study |
| sJMP12085 | A. baumannii AB5075 (sJMP3518) with attTn7::PabstBR-sfGFP (Tn7 from pJMP12049); aprR | this study |

[a]ampR, ampicillin resistant; aprR, apramycin resistant; kanR, kanamycin resistant; hygR, hygromycin resistant; dap-, requires diaminopimelic acid.

**Supplemental Table S2 Plasmids**

**Key**

| **Term** | **Description** |
|---|---|
| Plasmid # | JMP lab accession number |
| Description | Descriptive name |
| Addgene # | Addgene accession number (https://www.addgene.org/) |
| Construction/note | Plasmid construction notes, usage, or further details |
| Promoter-gene | Promoter used for expressing specific gene or reporter (if applicable) |
| Resistance | Antibiotic resistance cassette(s) |
| Source | plasmids produced in this study unless otherwise noted |

**Key plasmids from this study are shown in bold**

| Plasmid # | Description | Addgene # | Construction/notes | Promoter-gene | Resistance[a] | Source |
|---|---|---|---|---|---|---|
| pJMP0631 | Tiny Tn7 vector | | pTinyTn7 plasmid, R6k ori vector with Tn7 transposon with kanR cassette | na | ampR, kanR | Hall, PMID: 37662258 |
| pJMP1039 | pTn7C1 | 119239 | Tn7 transposase expression | na | ampR | Peters, PMID: 30617347 |
| pJMP2748 | vector that contains GFP | | sfGFP is on this vector for amplification | na | ampR, gentR | Ward, PMID: 38126769 |
| pJMP3067 | pTrc99a | | pTrc99a expression plasmid, pBR322 ori expression vector containing IPTG inducible trc promoter | Ptrc-empty | ampR | Rhodius, PMID:16336047 |
| pJMP3262 | pJQ200SK with p15A ori | 78497 | pJQ200SK, gent-sacB vector containing E. coli p15A origin of replication | na | gentR | Quandt, PMID: 8486283 |
| pJMP3341 | Tiny Tn7 with NcoI site removed | | Tiny-Tn7 plasmid; HiFi assembly to delete NcoI sites from pJMP631 with o1255/1258 and o1256/1257 amplified from pJMP631 | na | ampR, kanR | this study |
| pJMP3347 | pSGAb-km with pWH1266 ori | 121999 | plasmid containing pWH1266 A. baumannii origin of replication | na | kanR | Wang, PMID: 31548010 |
| pJMP3352 | Old Ptrc-EV expression vector, R6k version | | overexpression plasmid containing pTrc99a expression system, Ecoli R6K gamma origin, Abaumannii pWH1266 origin; (requires pir to replicate in E. coli) | Ptrc-empty | kanR | this study |
| pJMP3361 | Old Ptrc-GFP expression vector, R6k version | | overexpression plasmid containing Ptrc promoter expressing GFP, Ecoli R6K gamma origin, Abaumannii pWH1266 origin; HiFi assembled sfGFP gene amplified from pJMP2748 with oJMP1253/oMP1254 into pJMP3352 cut with NcoI/BamHI; (requires pir to replicate in E. coli) | Ptrc-GFP | kanR | this study |
| pJMP3407 | Old Pabst-EV expression vector, R6k version | | overexpression plasmid containing Pabst promoter, Ecoli R6K ori, Abaumannii pWH1266 ori; construced by PCR amplifying pJMP3352 with oJMP2040 and oJMP2041, and HiFi assembling with gblock oJMP2039 and pJMP3352 digested with EcoRI (used larger piece from digest via gel purification); (requires pir to replicate in E. coli) | Pabst-empty | kanR | this study |
| pJMP3434 | Old Pabst-GFP expression vector, R6k version | | overexpression plasmid containing Pabst promoter expressing GFP, Ecoli R6K gamma origin, Abaumannii pWH1266 origin; HiFi assembled sfGFP gene amplified from pJMP2748 with oJMP1253/oMP1254 into pJMP3407 cut with NcoI/BamHI; (requires pir to replicate in E. coli) | Pabst-GFP | kanR | this study |
| **pJMP3649** | **Ptrc EV kan - replicative vector used in this study** | | **Ptrc OE empty vector with p15A(ACYC) and pWH1266 origins; constructed by digesting pJMP3352 with AscI/PmeI, and HiFi assembling with PCR product of oJMP2134 and oJMP2135 amplified from pJQ200SK** | **Ptrc-empty** | **kanR** | **this study** |
| pJMP3650 | Ptrc GFP kan - replicative vector used in this study | | Ptrc-GFP expression vector with p15A(ACYC) and pWH1266 origins; constructed by digesting pJMP3361 with AscI/PmeI, and HiFi assembling with PCR product of oJMP2134 and oJMP2135 amplified from pJQ200SK | Ptrc-GFP | kanR | this study |
| **pJMP3651** | **Pabst EV kan - replicative vector used in this study** | | **Pabst OE empty vector with p15A(ACYC) and pWH1266 origins; constructed by digesting pJMP3407 with AscI/PmeI, and HiFi assembling with PCR product of oJMP2134 and oJMP2135 amplified from pJQ200SK** | **Pabst-empty** | **kanR** | **this study** |
| pJMP3652 | Pabst GFP kan - replicative vector used in this study | | Pabst-GFP expression vector with p15A(ACYC) and pWH1266 origins; constructed by digesting pJMP3434 with AscI/PmeI, and HiFi assembling with PCR product of oJMP2134 and oJMP2135 amplified from pJQ200SK | Pabst-GFP | kanR | this study |
| **pJMP3653** | **PabstBR EV kan - replicative vector used in this study** | * | **PabstBR OE empty vector with p15A(ACYC) and pWH1266 origins; constructed by digesting pJMP4481 with AscI/PmeI, and HiFi assembling with PCR product of oJMP2134 and oJMP2135 amplified from pJQ200SK** | **PabstBR-empty** | **kanR** | **this study** |
| **pJMP3654** | **PabstBR GFP kan - replicative vector used in this study** | * | **PabstBR-GFP epxression vector with p15A(ACYC) and pWH1266 origins; constructed by digesting pJMP4491 with AscI/PmeI, and HiFi assembling with PCR product of oJMP2134 and oJMP2135 amplified from pJQ200SK** | **PabstBR-GFP** | **kanR** | **this study** |
| **pJMP3664** | **PabstBR EV apr - replicative vector used in this study** | * | **Apramycin version of pJMP3653 (PabstBR-empty vector); digested pJMP3653 with XhoI, HiFi assembled with gblock oJMP1946** | **PabstBR-empty** | **aprR** | **this study** |
| **pJMP3665** | **PabstBR EV hyg - replicative vector used in this study** | * | **Hygromycin version of pJMP3653 (PabstBR-empty vector); digested pJMP3653 with XhoI, HiFi assembled with gblock oJMP2117** | **PabstBR-GFP** | **hygR** | **this study** |
| pJMP3666 | PabstBR GFP apr - replicative vector used in this study | * | Apramycin version of pJMP3654 (PabstBR-empty vector); digested pJMP3654 with XhoI, HiFi assembled with gblock oJMP1946 | PabstBR-empty | aprR | this study |
| pJMP3667 | PabstBR GFP hyg - replicative vector used in this study | * | Hygromycin version of pJMP3654 (PabstBR-empty vector); digested pJMP3654 with XhoI, HiFi assembled with gblock oJMP2117 | PabstBR-GFP | hygR | this study |
| pJMP3748 | Tn7 PrpoE-mRFP reporter vector | | Integrative expression vector with PrpoE-mRFP reporter on Tn7 transposon; derived from pJMP8602 by adding PrpoE in front of mRFP reporter using oligos oJMP2369 and oJMP2370 | PrpoE-mRFP | ampR, aprR | this study |
| pJMP3853 | PabstBR rpoE OE vector kan | | PabstBR-RpoE expression vector, parent pJMP3653; HiFi assembled rpoE gene amplified from E. coli sJMP3075 gDNA with oJMP2634 and oJMP2635 into pJMP3653 digested with NcoI/BamHI | PabstBR-RpoE | kanR | this study |
| pJMP4003 | pEVS104 helper plasmid | 207393 | pEVS104(R6K ori, KanR, Tra+) | na | kanR | Stabb, PMID: 12474404 |
| pJMP4481 | Old PabstBR-EV expression vector, R6k version | | Overexpression plasmid containing PabstBR promoter, Ecoli R6K ori, Abaumannii pWH1266 ori; construced by site-directed mutagenesis using pJMP3407 and oJMP2167; (requires pir to replicate in E. coli) | PabstBR-empty | kanR | this study |
| pJMP4491 | Old PabstBR-GFP expression vector, R6k version | | Overexpression plasmid containing PabstBR promoter expressing GFP, Ecoli R6K gamma origin, Abaumannii pWH1266 origin; HiFi assembled sfGFP gene amplified from pJMP2748 with oJMP1253/oMP1254 into pJMP4481 cut with NcoI/BamHI; (requires pir to replicate in E. coli) | PabstBR-GFP | kanR | this study |
| pJMP8602 | Old vector containing Tn7 base | | Tn7 expression vector with mRFP reporter on Tn7 transposon | none-mRFP | kanR | Hall, PMID: 37662258 |
| **pJMP12042** | **Tn7 PabstBR EV apr - integrative vector used in this study** | * | **Integrative expression vector with PabstBR empty vector on Tn7 transposon; constructed by amplifying pJMP3653 with oJMP2539 and oJMP2540, pJMP8602 with oJMP2541 and oJMP2542, pJMP3352 with oJMP2543 and oJMP2544, and HiFi assembling with pJMP3539 digested with SpeI/NotI** | **PabstBR-empty** | **ampR, aprR** | **this study** |
| **pJMP12049** | **Tn7 PabstBR GFP apr - integrative vector used in this study** | * | **Integrative expression vector with PabstBR-GFP on Tn7 transposon; constructed by digesting pJMP12042 with NcoI/BamHI, and HiFi assembling with PCR product of oJMP1254 and oJMP2078 amplified from pJMP2748** | **PabstBR-GFP** | **ampR, aprR** | **this study** |
| **pJMP12068** | **Tn7 PabstBR EV kan - integrative vector used in this study** | * | **Kanamycin version of pJMP12042 (Tn7-PabstBR-empty vector); HiFi assembled PCR product of pJMP3653 with oJMP0193 and oJMP0194 into pJMP12042 digested with XhoI** | **PabstBR-empty** | **ampR, kanR** | **this study** |
| **pJMP12089** | **Tn7 PabstBR EV hyg - integrative vector used in this study** | * | **Hygromycin version of pJMP12042 (Tn7-PabstBR-empty vector); digested pJMP12042 with XhoI, HiFi assembled with gblock oJMP2117** | **PabstBR-empty** | **ampR, hygR** | **this study** |

[a]ampR, ampicillin resistant; aprR, apramycin resistant; gentR, gentamycin resistant; kanR, kanamycin resistant

*to be submitted to Addgene

**Supplemental Table S3 Oligonucleotides and Synthetic DNA**

**Key**

| **Term** | **Description** |
|---|---|
| Oligo # | JMP lab accession number |
| Sequence | Nucleotide sequence |
| Usage | Procedure using this oligonucleotide/synthetic DNA |

| Oligo # | Sequence[a] | Usage |
|---|---|---|
| oJMP0193 | TCCTGAACGGCCATAAGAAC | amplifying kanR marker for swapping A |
| oJMP0194 | GAGCGCTTTTGAAGCTGATG | amplifying kanR marker for swapping B |
| oJMP1253 | CGGATAACAATTTCACACAGGAAACAGACCATGAGCAAAGGAGAAGAACT | amplify GFP A |
| oJMP1254 | TGCATGCCTGCAGGTCGACTCTAGAGGATCTTTGTAGAGCTCATCCATGC | amplify GFP B |
| oJMP1255 | TTCGGCAAGCAGGCATCGCCGTGGGTCACGACGAGATCCT | remove NcoI sites from pJMP631 A |
| oJMP1256 | GAGGATCTCGTCGTGACCCACGGCGATGCCTGCTTGCCGA | remove NcoI sites from pJMP631 C |
| oJMP1257 | CTGACATGGGAATTAGCCACGGCATCACAGTATCGTGATG | remove NcoI sites from pJMP631 D |
| oJMP1258 | ATCACGATACTGTGATGCCGTGGCTAATTCCCATGTCAGC | remove NcoI sites from pJMP631 B |
| oJMP1946 | TCCTGAACGGCCATAAGAACGAAGGCTGTCTGTTGAACTCTCGAGCCTTGTCGCCTTGCGTATAATATTTGCTCA | apramycin (aprR) gblock |
| oJMP2039 | AGCCATCGGAAGCTGTGGTATGGCTGTGCAGGTCGTAAAGACGTCTCATTAATCAGATAAAATATTATAAATGTC | making Pabst replicative vector gblock |
| oJMP2040 | TGATACCTCTGGCTCAGCG | making Pabst replicative vector A |
| oJMP2041 | TTTACGACCTGCACAGCCAT | making Pabst replicative vector B |
| oJMP2117 | TCCTGAACGGCCATAAGAACGAAGGCTGTCTGTTGAACTCTCGAGCCTTGTCGCCTTGCGTATAATATTTGCTCA | hygromycin (hygR) gblock |
| oJMP2134 | GCCTTCCGTTTGGGCCTAGAATTCCGGTTTAACAGGGAAGTGAGAGGGC | swapping to p15A ori A |
| oJMP2135 | TGGCTTGCAGCCCTTATGTCCTCAGGCGCGCACAAAACAGCAGGGAAGC | swapping to p15A ori B |
| oJMP2167 | CAAGAAGATCATCTTATTAATCAGATAAAATATTGACAATGTGAGCGGATAACAAG | making PabstBR Quikchange primer |
| oJMP2369 | CTAGAAACAAAAACAGATGCGTTACGGAACTTTACAAAAACGAGACACTCTAACCCTTTG | PrpoE cloning A |
| oJMP2370 | TCGTCAAAGGGTTAGAGTGTCTCGTTTTTGTAAAGTTCCGTAACGCATCTGTTTTTGTTT | PrpoE cloning B |
| oJMP2539 | CACGCCGCTTTTTTTACGTCTGCAGACTAGTCAGGCAGCCATCGGAAGCT | amplify MCS A |
| oJMP2540 | TGTCGACTAGGCCATGATGCTCATTCTGTGAAAAGGCCATCCGTCAGGAT | amplify MCS B |
| oJMP2541 | CACAGAATGAGCATCATGGC | amplify Tn7 cassette A |
| oJMP2542 | GGGTCAGTGAGCGAGGAAG | amplify Tn7 cassette B |
| oJMP2543 | TTTAAAACTTCATTTTTAATTTTTGCGGCCGATCTGAAGATCAGCAGTTC | amplify R6k and oriT A |
| oJMP2544 | GGGCTCTTCCGCTTCCTCGCTCACTGACCCTATCAGAGCTTATCGGCCAG | amplify R6k and oriT B |

[a]all sequences are 5'-3'
